# A chemical interpretation of protein electron density maps in the worldwide protein data bank

**DOI:** 10.1101/613109

**Authors:** Sen Yao, Hunter N.B. Moseley

## Abstract

High-quality three-dimensional structural data is of great value for the functional interpretation of biomacromolecules, especially proteins; however, structural quality varies greatly across the entries in the worldwide Protein Data Bank (wwPDB). Since 2008, the wwPDB has required the inclusion of structure factors with the deposition of x-ray crystallographic structures to support the independent evaluation of structures with respect to the underlying experimental data used to derive those structures. However, interpreting the discrepancies between the structural model and its underlying electron density data is difficult, since derived electron density maps use arbitrary electron density units which are inconsistent between maps from different wwPDB entries. Therefore, we have developed a method that converts electron density values into units of electrons. With this conversion, we have developed new methods that can evaluate specific regions of an x-ray crystallographic structure with respect to a physicochemical interpretation of its corresponding electron density map. We have systematically compared all deposited x-ray crystallographic protein models in the wwPDB with their underlying electron density maps, if available, and characterized the electron density in terms of expected numbers of electrons based on the structural model. The methods generated coherent evaluation metrics throughout all PDB entries with associated electron density data, which are consistent with visualization software that would normally be used for manual quality assessment. To our knowledge, this is the first attempt to derive units of electrons directly from electron density maps without the aid of the underlying structure factors. These new metrics are biochemically-informative and can be extremely useful for filtering out low-quality structural regions from inclusion into systematic analyses that span large numbers of PDB entries. Furthermore, these new metrics will improve the ability of non-crystallographers to evaluate regions of interest within PDB entries, since only the PDB structure and the associated electron density maps are needed. These new methods are available as a well-documented Python package on GitHub and the Python Package Index under a modified Clear BSD open source license.

**Author summary:** Electron density maps are very useful for validating the x-ray structure models in the Protein Data Bank (PDB). However, it is often daunting for non-crystallographers to use electron density maps, as it requires a lot of prior knowledge. This study provides methods that can infer chemical information solely from the electron density maps available from the PDB to interpret the electron density and electron density discrepancy values in terms of units of electrons. It also provides methods to evaluate regions of interest in terms of the number of missing or excessing electrons, so that a broader audience, such as biologists or bioinformaticians, can also make better use of the electron density information available in the PDB, especially for quality control purposes.

**Software and full results available at:** https://github.com/MoseleyBioinformaticsLab/pdb_eda (software on GitHub)

https://pypi.org/project/pdb-eda/ (software on PyPI)

https://pdb-eda.readthedocs.io/en/latest/ (documentation on ReadTheDocs)

https://doi.org/10.6084/m9.figshare.7994294 (code and results on FigShare)

## Introduction

Proteins are active components in the biochemical implementation of biological processes, and understanding their structure is important for interpreting their biochemical functions. The Worldwide Protein Data Bank (wwPDB, www.wwpdb.org) [1] is the international organization that manages the Protein Data Bank (PDB, www.rcsb.org) [2], the central repository of biological macromolecules structures. Thousands of structures are deposited into the wwPDB every year, but their data quality can vary significantly from structure to structure, and even region to region within a structure. Low-quality data can cause problems for both a single macromolecule structure inspection and aggregated systematic analyses across hundreds or thousands of structural entries [3, 4]. Thus, the analysis and interpretation issues caused by the presence of low-quality structural data are pushing the structural biology community to pay more attention to the quality of deposited structural entries [5]. The wwPDB has initiated several efforts to improve the quality of entries being deposited, including launching a deposition, biocuration, and validation tool: OneDep [6]. Many data quality measures are now available for PDB structures, such as a resolution, b-factors, MolProbity clashscores [7], to name a few. However, low-quality regions can still exist even in structures with very good metrics of global structural quality, as shown in the overlay of structures with electron density maps in Fig 1. These low-quality regions arise from structural model and electron density mismatches can be due to a variety of reasons including problems with regional protein mobility [8], data processing [9], or model fitting [10]. These mismatches often occur around bound ligands where a lot of interesting biological activities happens, making the analysis of protein sequence-structure-function relationships more difficult. Therefore, evaluation of structure quality, especially around regions of interest, is paramount before accurate structural inferences can be made.

**Fig 1.**
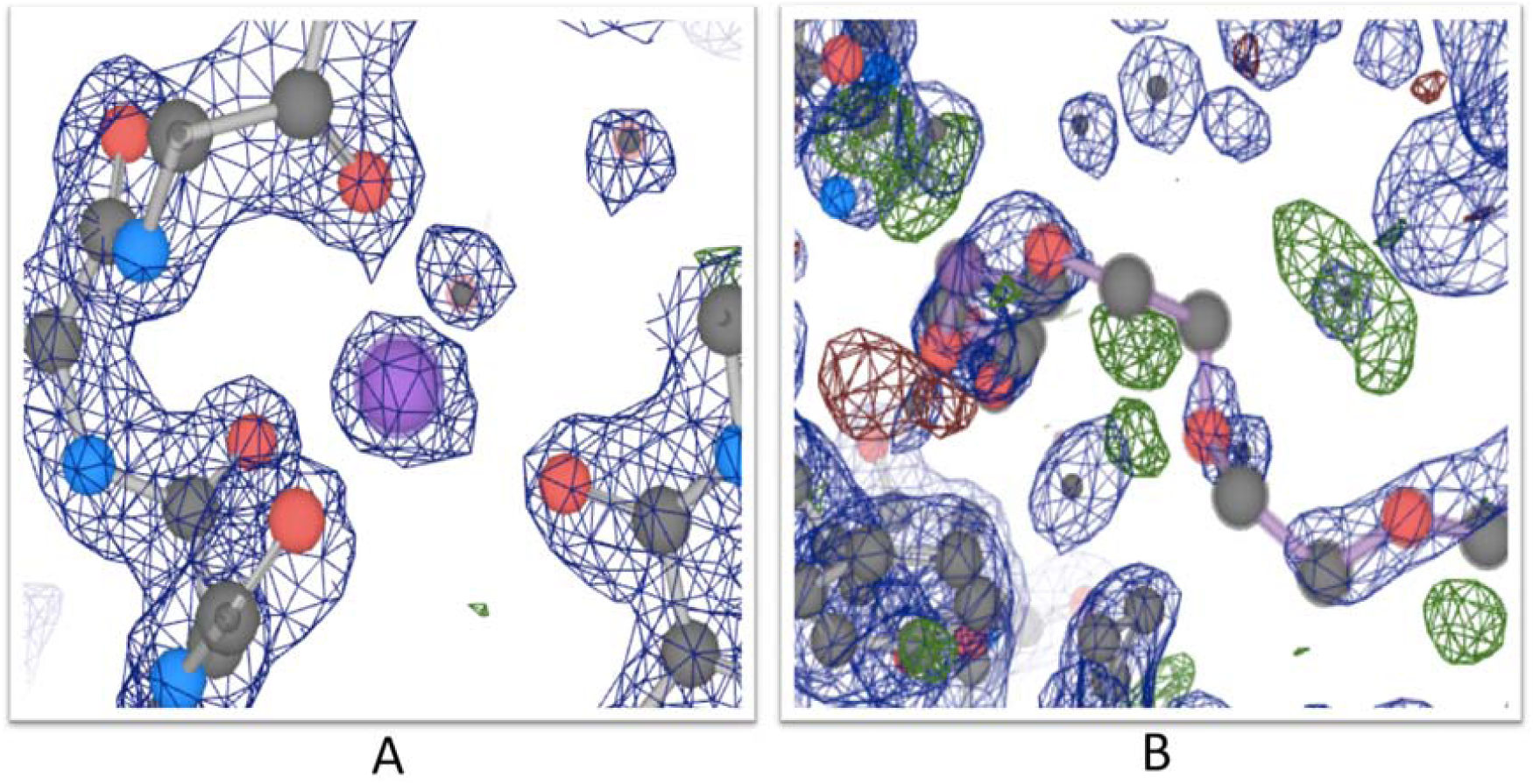
Model and difference electron density maps. Both panels A and B are from the same structure, PDB ID: 3B1Q, and are centered around the ligands B.330 (A) and P.33 (B). Blue meshes represent the model electron density. Green and red meshes represent discrepancies between experimental data and the structure model. The structure has very good overall quality, as demonstrated by a resolution of 1.7 Å, an R-factor value of 0.176, and an R-free value of 0.207. The panel A shows a high-quality region in the structure where the experimental data and model match very well. And the panel B shows that there are still low-quality regions within the structure, as demonstrated by the green and red blobs around the coordinating ligand residue (“ligand” refers to the coordination chemistry definition of this word).

Driving improved evaluations of structure quality are newer deposition requirements like mandatory deposition of structure factors (x-ray structures) and constraints (NMR structures) starting from 2008 [11], NMR-assigned chemical shifts from 2010, and 3DEM volume maps from 2016 [7]. The inclusion of underlying experimental data used in structure determination enables researchers to better validate structural models, improving the inferences they can make from these structures. For x-ray crystallographic structures, electron density maps enable a direct comparison between the observed electron density Fo to calculated electron density Fc based on the structural model. The 2Fo-Fc map represents the electron densities surrounding well-determined atoms in the model across a three-dimensional (3D) space and the Fo-Fc map represents the electron density discrepancies between the observed and calculated electron density across a 3D space. For x-ray crystallographic PDB entries with deposited experimental data, electron density maps are made available by the PDB in Europe (PDBe) [12]. Previously from about 1998 to 2018 [13], electron density maps were made available by the Uppsala Electron Density Server (Uppsala EDS), which was created and maintained outside of the PDB [14]. However, the PDBe uses newer methods to provides higher quality density maps, which prompted the retirement of the Uppsala EDS by 2018. Many electron density map viewers [15, 16] exist for manually examining the quality of a model versus its electron density; however, this software and evaluation approach is not suitable for batch analysis of hundreds of structures. Also, these electron density maps are in arbitrary units of electron density, with no direct physicochemical meaning. This normally does not affect the visualization of electron densities and is a by-product of creating maps with a summative intensity of zero (zero-sum) across the whole map, which is done primarily for visual simplification during modeling [17, 18]. But this zero-sum representation can be detrimental for understanding a model, especially a local region of a model, where the number of electrons of density or density discrepancy would be useful for evaluation. Due to these limitations, we have developed a new method that derives a conversion factor from the arbitrary electron density units of a given electron density map with corresponding PDB entry into the absolute value of electrons per angstroms cubed. With this conversion factor, we have developed new evaluation methods that normalize electron density and electron density discrepancies into estimated quantities of electrons. These new electron discrepancy values can provide chemically-informative information for evaluating structural models or for filtering structure entry regions for inclusion into systematic analyses that span large numbers of PDB entries.

## Methods

### Calculating the electron density ratio for atoms, residues, and chains

A workflow of the analysis is shown in Fig 2. Structural data was downloaded from wwPDB on Jul 3, 2018, and their electron density data, if available, was acquired from the PDBe website [12]. Structural data was processed using a self-developed parser and Biopython [19]. Electron density data was analyzed according to the CCP4 suite [20] format guidance. The electron density map is represented as a 3D array in the data, which corresponds to voxels in the real space. An electron density voxel with a density value greater than 1.5σ of all voxels is considered significant for 2Fo-Fc maps, and 3σ for Fo-Fc maps. For structures with electron density data available, symmetry operations were performed to include the surrounding environment for the modeled structure.

**Fig 2.**
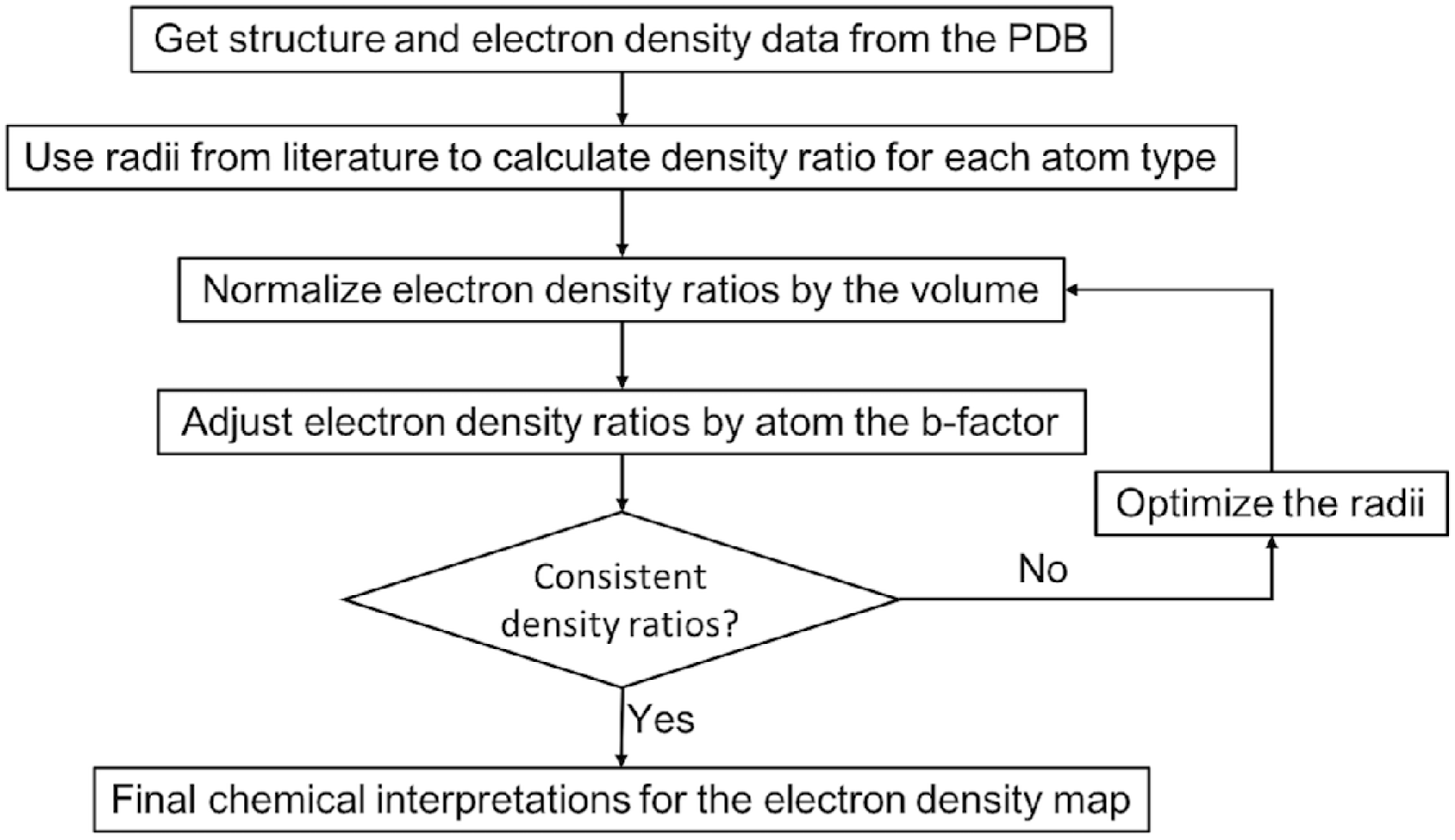
Workflow of the electron density analysis.

To calculate the total electron density around each atom, we initially used the radii from literature [21] and calculated the sum of all densities within the corresponding radius. The voxel center is used when calculating the distance from a density voxel to an atom. Different atoms (without hydrogens) from the 20 common amino acid are categorized into 13 atom types as shown in Table S1. Electron density ratio (*r*_*i*_) is defined as the total density of all associated voxels (ρ_m_) divided by the number of electrons (Z_i_) for a given atom *i*,

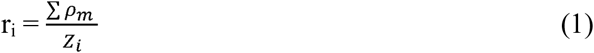

As the unit for electron density is eÅ^-3^, the electron density ratio thus has the unit as Å^-3^. The total density is adjusted by a factor of the occupancy of the atom. Since hydrogen is normally not resolvable within electron density maps, their electrons were added to their bonded atom. A table of electron counts used for each atom is shown in Table S2.

After all atom electron densities are calculated, they are aggregated into residue and chain densities where the residue cloud contains at least 4 atoms and the chain cloud contains at least 50 atoms. The overlapping density voxels between two or more atoms are only counted once through the aggregation. The total number of electrons are calculated by adding contributing atom’s electron numbers together. Residue (*r*_*r*_) and chain (*r*_*c*_) density ratios are then calculated accordingly.

### Normalizing the electron density ratio by the number of voxels

To smooth the representation of continuous electron densities using discrete voxels, the electron density ratio is then normalized by the median volume (in number of voxels) of a given atom type. If we denote the original density ratio as *r*_*i*_ and the volume of a given atom *i* with atom type *t* as *V*_*i*_, and the median volume of all atoms with atom type *t* as *median(V*_*t*_), the normalized density ratio *r*_*i-norm*_ can be defined as follow:

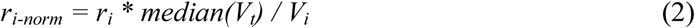

### Correcting the unit electron density by the atom b-factor

As the actual value of the density ratio is highly specific to individual structures, we then define a more universal measure as the chain deviation fraction (*f*_*i*_) for a given atom *i* as:

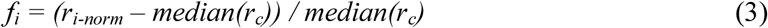

The dispersion of electron density around an atom can be approximated using the b-factor of the given atom. The chain fraction and logarithmic b-factor have a linear correlation both statistically and visually, and thus a slope (*s*_*t*_) of chain fraction over logarithmic b-factor can be calculated for each atom type and for every structure. If there are less than three points for a given atom type, the median slope over 1000 random structures is used. Then for each individual atom *i* with atom type *t*, its unit electron density can be corrected by its deviation from the median b-factor:

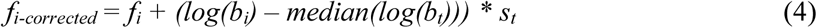

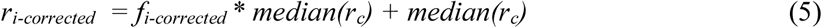

### Optimization of radii

After the initial calculation, the median density ratios of different atom types were still quite different from each other. Thus, to achieve a more uniformly interpretable density ratio within a structure as well as across structures, an optimization of radii is performed. First, we tested the radius for each atom type on 100 random structures and obtained an initial estimation of the radii. The metric we used to optimize is the median of corrected chain deviation fraction (*f*_*i-corrected*_) for a given atom type. Based on the results from the initial step, we then optimize one atom type at a time on 1000 randomly selected structures. For every iteration, the atom type that has the largest deviation from last round is optimized. Different radii are tested for the given atom type and the radius that has a median corrected chain deviation fraction closest to zero is picked out. At the end of each optimization, the set of b-factor slopes are updated as well. This process goes on until the median chain deviation fraction for all atom types are smaller than 0.05. The final set of radii are then tested on another 1000 random structures and the whole PDB database for validation.

### Adding an F_000_ term

The average value of the 2mFo-DFc map (i.e. a Sigma-A weighted map) is practically zero for most of the structures in the PDB. Theoretically, an F_000_ term should be added to get the proper number of electrons on an absolute scale. Unfortunately, not all structure factor programs provide the F_000_ value. So as an estimation, we add up the numbers of electrons for all the atoms of a model in the unit cell, including symmetry structure units and modeled water molecules, and then divide it by the number of voxels within the unit cell,

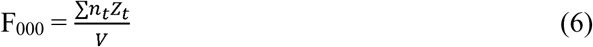

Where n_t_ is number of atoms of element t in the asymmetric unit, Z_t_ is the number of electrons (atomic number) of element t, and V is the unit cell volume in Å^3^. This estimated F_000_ term is then added to the density values of all the voxels.

### Design, implementation, and distribution of the above methods

These new methods are implemented in a Python package, pdb-eda. It is written in major version 3 of the Python program language and is available on GitHub, https://github.com/MoseleyBioinformaticsLab/pdb_eda, and the Python Package Index (PyPI), https://pypi.org/project/pdb-eda/. There are three main parts of pdb-eda: the pdb parser, the ccp4 parser, and the electron density analysis. Starting from a PDB id, pdb-eda can either read a local pdb or ccp4 file or download it on the fly. Intermediate and final results of all three parts can be accessed via either importing as a library or using the command line interface. Many options are available for handling and processing the data with the details documented in the package guide and tutorial files. As part of the development process, new versions are updated on GitHub regularly. The version that is described and implemented for this paper has been frozen, tarballed, and published on FigShare (https://doi.org/10.6084/m9.figshare.7994294), along with all the result files and codes in generating all results, figures, and tables.

## Results

We downloaded and used a total of 141,763 wwPDB entries, of which 106,321 structures have electron density maps available and suitable for the analysis in this study. The assumption of this study is based on a fundamental rule for electron density construction, that is the electron density is proportional to the number of electrons. However, after being deposited into the PDB, this information of absolute value of electrons is hard to derive and is inconsistent across structures. Different structures can vary a lot in terms of density ratios, due to the quality of the crystal or the choice of data processing software. Therefore, we need an internal measure to enable a consistent interpretation within and across structures. If we simply use the radii from the literature and do not apply any correction, the median of atom density ratio shows that the density ratios are inconsistent within a single structure for atoms, residues, and chains, as illustrated in Fig 3 Panels A-C. The atom density ratios span over the largest range, while the chain density ratio has the smallest range. Therefore, we chose the chain deviation fraction as a reliable measure to optimize all atom types to the same level.

**Fig 3.**
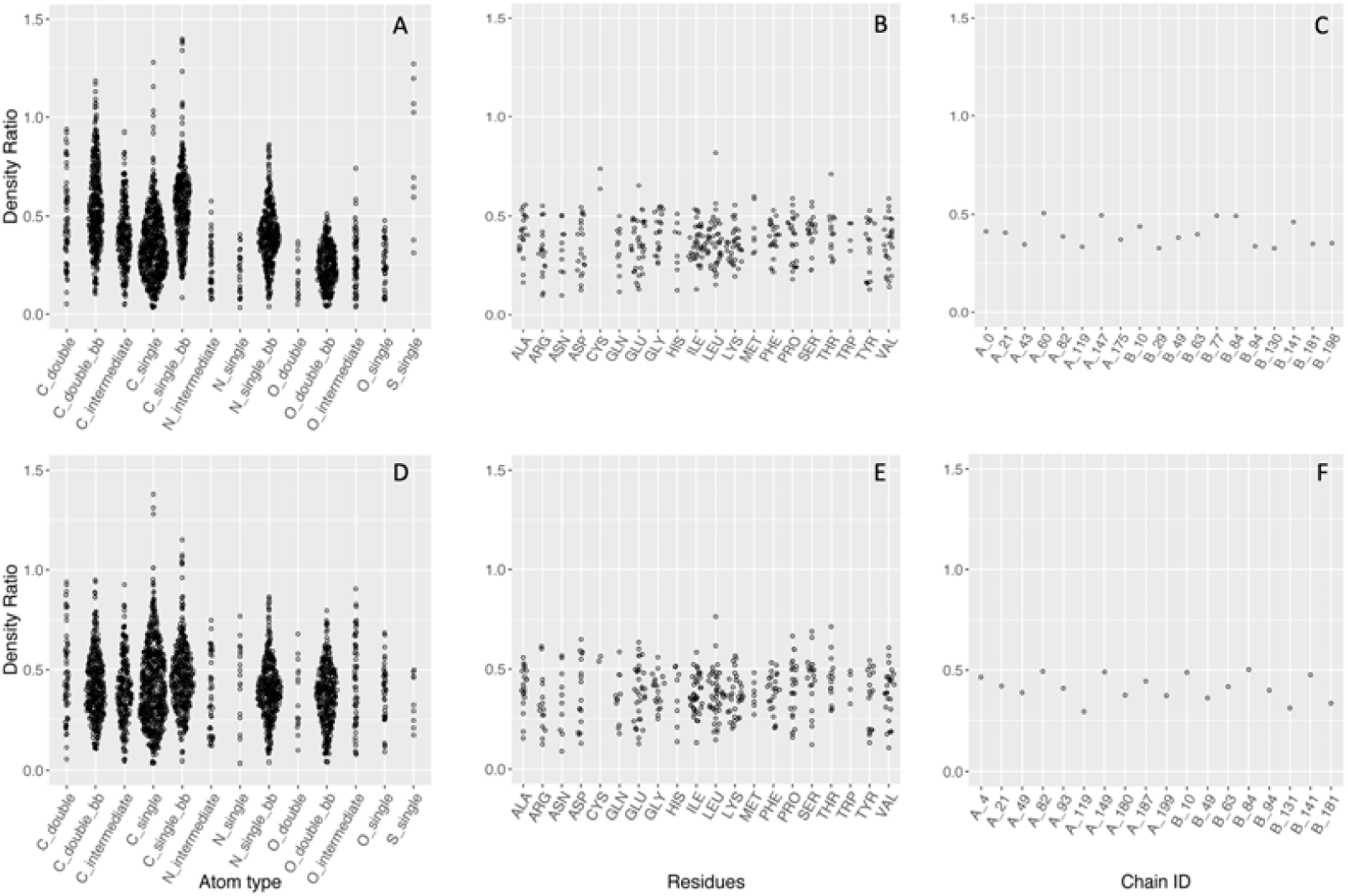
Sina plots of density ratio for atoms, residues, and chains, before (Panels A-C) and after (Panels D-F) radii optimization. PDB ID: 3UBK. The atom density ratios have the largest range, and chain density ratios have the smallest range, which can be used as internal standard to optimize the atom ratios to.

### Volume and b-factor adjustment

An example of the volume distribution for a single structure is shown in Fig 4. Ideally, atoms of the same atom type should occupy the same volume. However in reality, different parts of a structure may be more or less ordered within the crystal than others. Also, during the reconstruction of the electron density data from complex structure factors, continuous electron density through space is represented as a point density value for every voxel. And the use of the point density to estimate the whole voxel depends on the smoothness of the density function. Moreover, based on the placement of the atom in relation to the voxel and the selection of grid length, the inclusion of a voxel is an all-or-none decision. Thus, we performed the volume normalization to minimize these effects.

**Figure 4.**
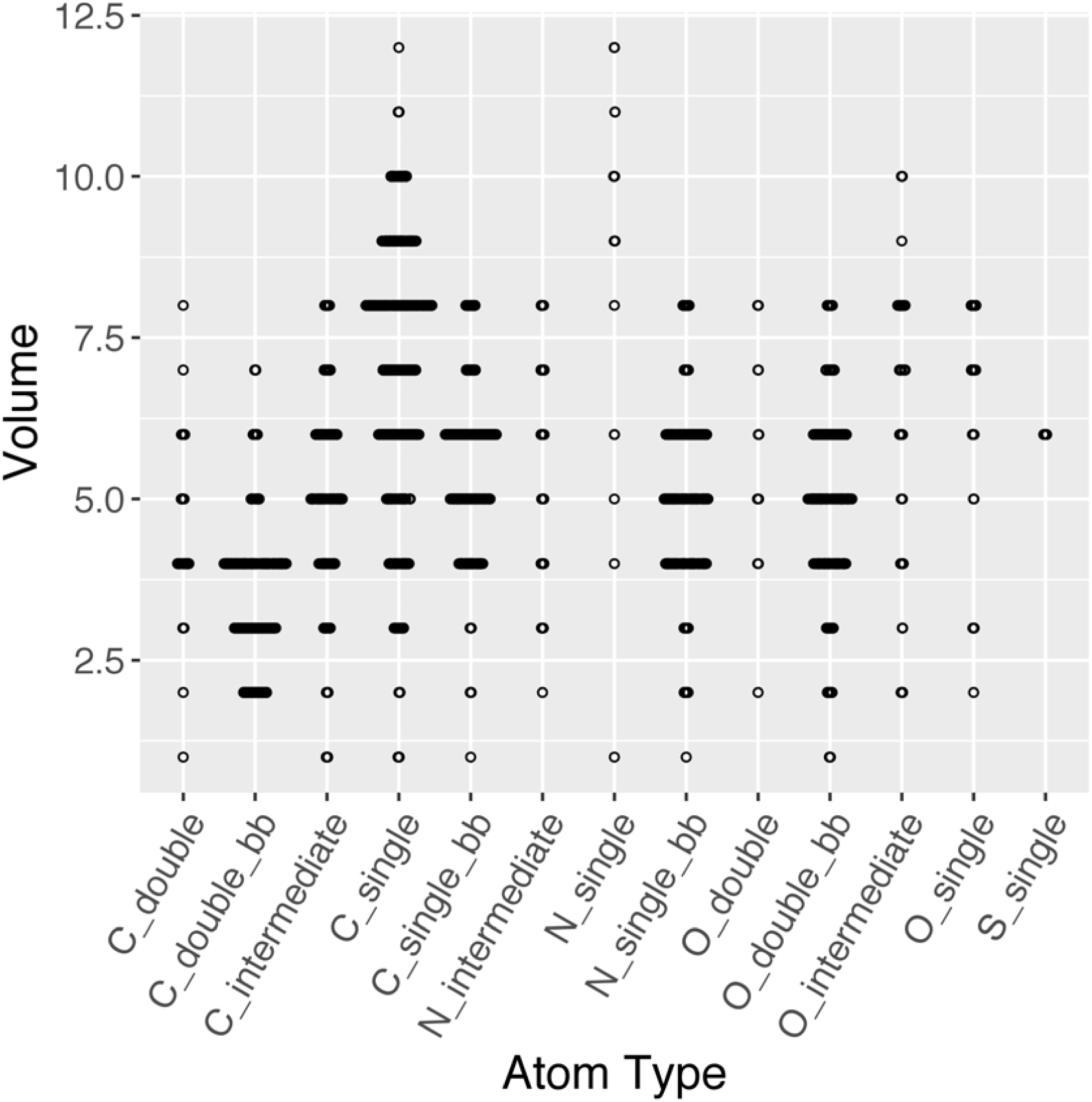
Sina plot for the volumes of each atom type. PDB ID: 3UBK.

B-factor measures the temperature-dependent atomic displacement in a crystal. As a result, it is inversely correlated with the total electron density within a distance of an atom, and thus the density ratio. After examining several different relationships between density ratio and b-factor, both statistically and visually, the chain deviation fraction versus logarithmic b-factor demonstrated the strongest linear correlation, and thus was used for the correction. Fig 5 provides an example of this relationship.

**Fig 5.**
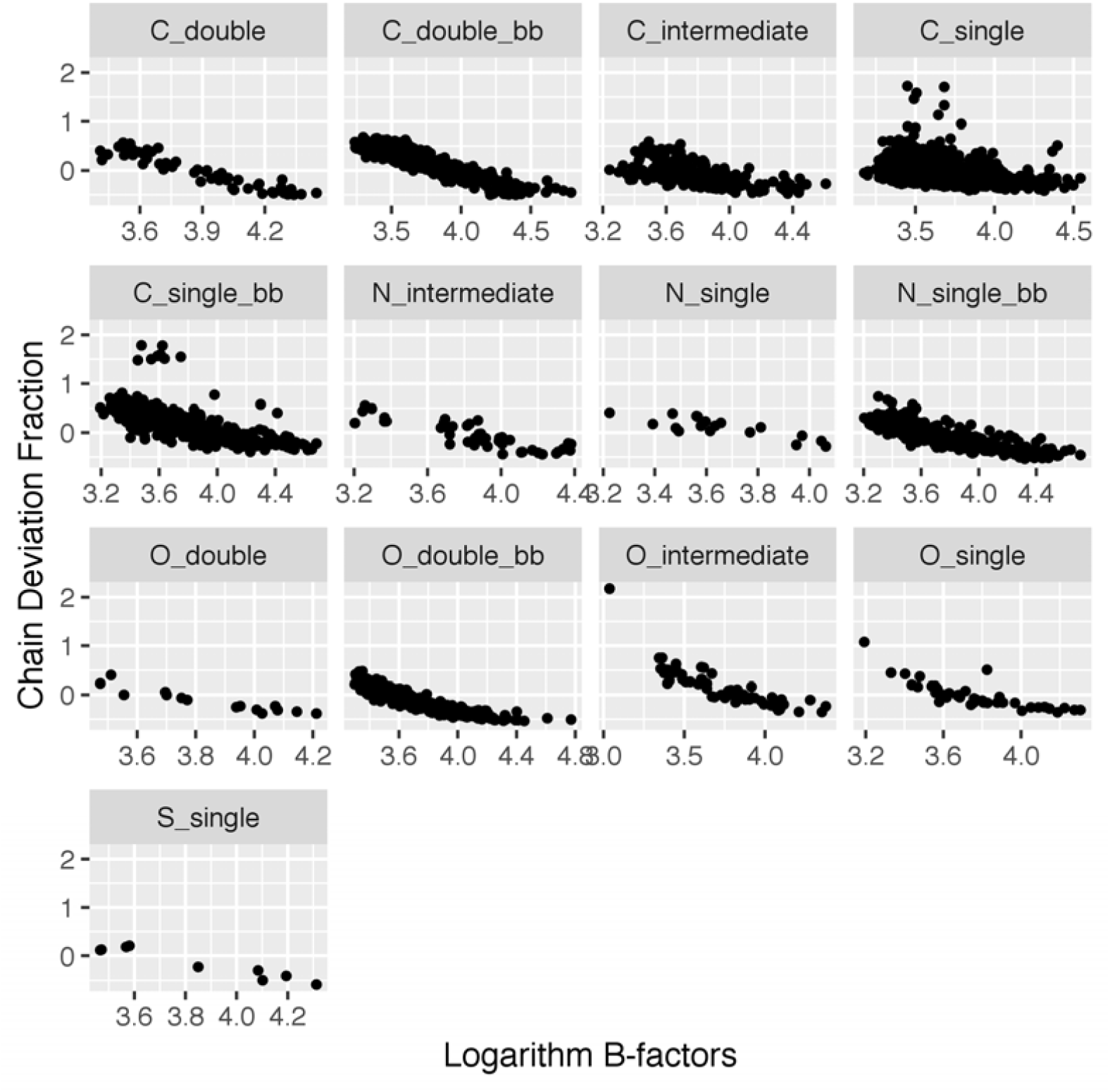
Correlation between chain deviation fractions and logarithm B-factors for each atom type. PDB ID: 3UBK.

Both volume normalization and b-factor correction help to reduce the high variability of the density ratio. Fig 6 illustrates the distributions of atom density ratios before and after each step (Panels A-C). After both adjustments, the atom density ratio is coherent within each atom type, though it is still uneven between atom types (Panel C).

**Fig 6.**
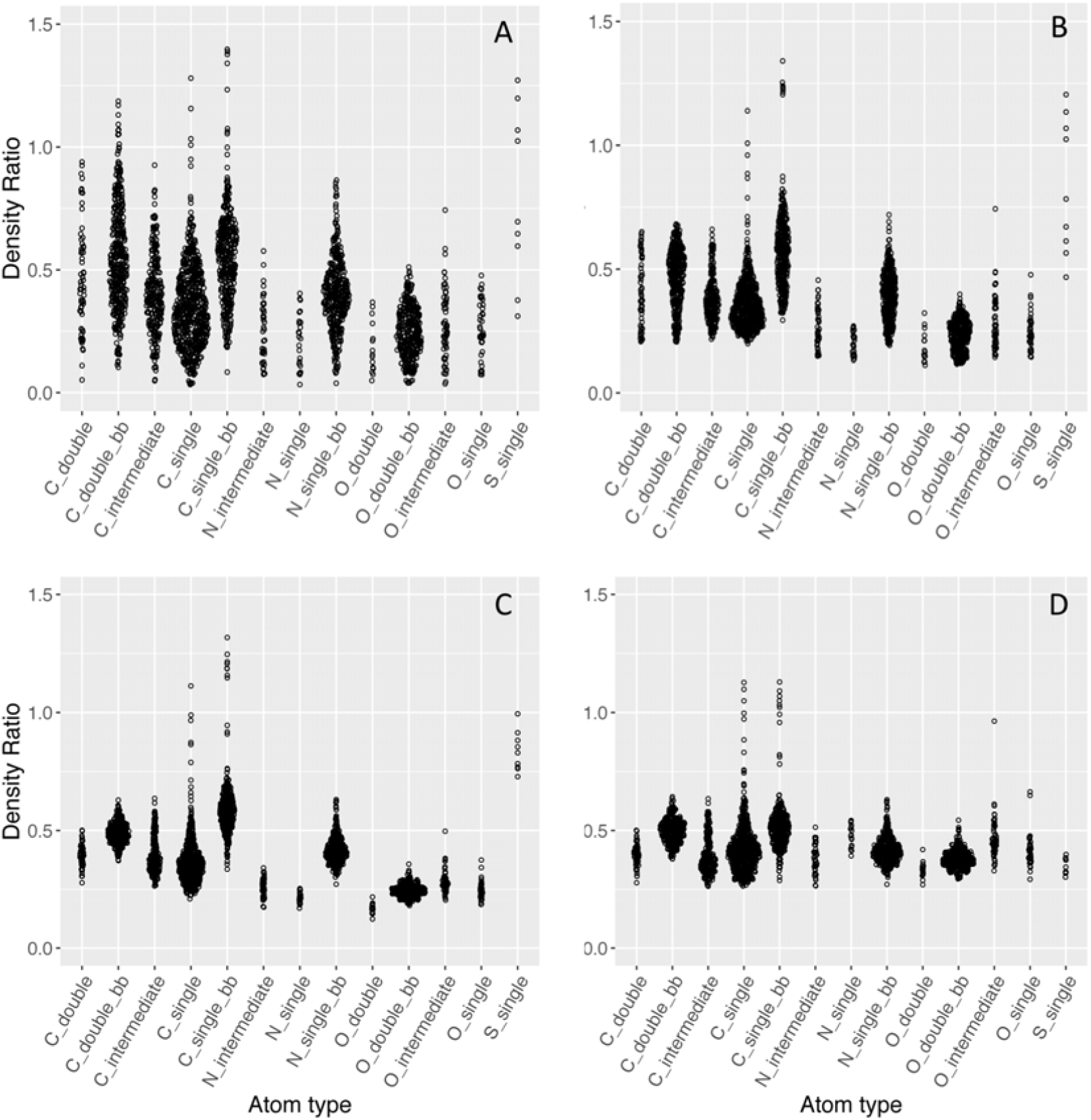
Atom density ratios at each major step of improvement. A) Original, B) After volume normalization, C) After b-factor correction, D) After radii optimization. PDB ID: 3UBK. After each major step, the distribution of the atom density ratios becomes less spread.

### The final set of radii after optimization

Fig 6, Panel D illustrates the distributions of atom density ratios after radii optimization, where different atom types have much more similar median ratios. A comparison of initial and final set of radii is shown in Table 1. In general, most of the backbone atoms decrease in radius, while most of the optimized radii on the side chain are larger than those on the backbone. This is due to the lower order and higher flexibility of the side chain atoms, which practically requires a larger radius to capture the expected number of electrons.

**Table 1.** The atom radii before and after radii optimization.

| Atom Type | Original Radius (Å) | Optimized Radius (Å) |
| --- | --- | --- |
| C_single | 0.77 | 0.84 |
| C_single_bb | 0.77 | 0.72 |
| C_double | 0.67 | 0.67 |
| C_double_bb | 0.67 | 0.61 |
| C_intermediate | 0.72 | 0.72 |
| O_single | 0.67 | 0.80 |
| O_double | 0.60 | 0.71 |
| O_double_bb | 0.60 | 0.77 |
| O_intermediate | 0.64 | 0.71 |
| N_single | 0.70 | 0.95 |
| N_single_bb | 0.70 | 0.70 |
| N_intermediate | 0.62 | 0.77 |
| S_single | 1.04 | 0.75 |

The radius of sulfur changes the most as compared to the other elements. This could be due to how most software construct electron density data from structure factor and associated phase via Fourier transforms. The electron density is approximated with a Gaussian distribution, and its variance is affected mainly by the b-factor. Thus, the final optimized radius is a combination of the actual radius of the atom, the displacement of an atom center, as well as the thermal motion of the atom (B factor). And studying the behavior of sulfur atoms could be useful for other less common elements such as metal ions.

### Overview of density ratio for the whole PDB database

The final set of radii was first tested on another 1000 random structures and all atom types hold true to have no more than a 5% chain deviation fraction. It was then applied to the all PDB structures that has usable electron density data, and the results are shown in Fig 7. For all atom types the distributions center around 0, which indicates the set of optimized radii yields consistent measures of density ratios across structures. As shown in Fig S1, for high-quality structures with a resolution smaller than 1.5Å, the chain deviation fraction illustrates tighter distributions with modes above 0, because of the narrower electron dispersion around atoms in the experimental data. As the resolution gets worse, this distribution tends to broaden for all atom types with the modes smaller than 0.

**Fig 7.**
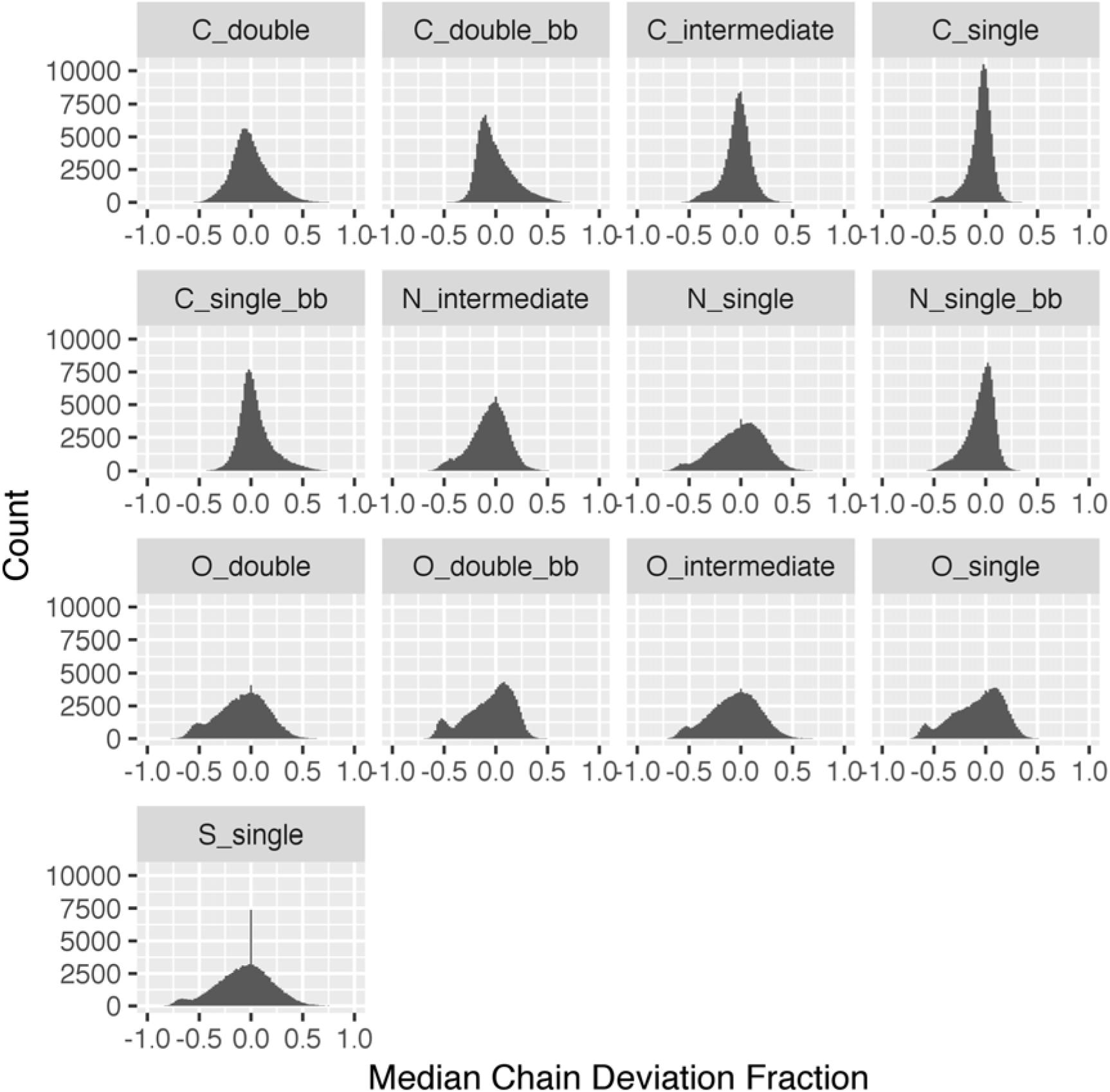
Histogram of the median chain deviation fraction for all structures in the PDB.

### F_000_ term and the absolute scale

Analysis of electron density on an absolute scale (i.e. in units of e/Å^3^) requires the value of F_000_ and the unit cell volume. As the shape of the density matters more than the absolute scale in the structure modeling, most maps lack this F_000_ term. Therefore, the mean value of the 2mFo-DFc map is practically zero across the whole PDB, as shown in Fig 8. To get the true electron density values, we would need to add an F_000_ term to the set of Fourier coefficients going into the calculation of the map. However, as Fig 9 shows before and after adding the estimated F_000_ term, it makes very little contribution to the overall absolute electron density values. Thus, conversion is not as simple as adding an F_000_ term as often theoretically represented in textbooks [22, 23] and likely depends on software parameters used in the creation of the map. Therefore, the chain median is used as a conversion factor to relate all electron density values back to the absolute scale.

**Fig 8.**
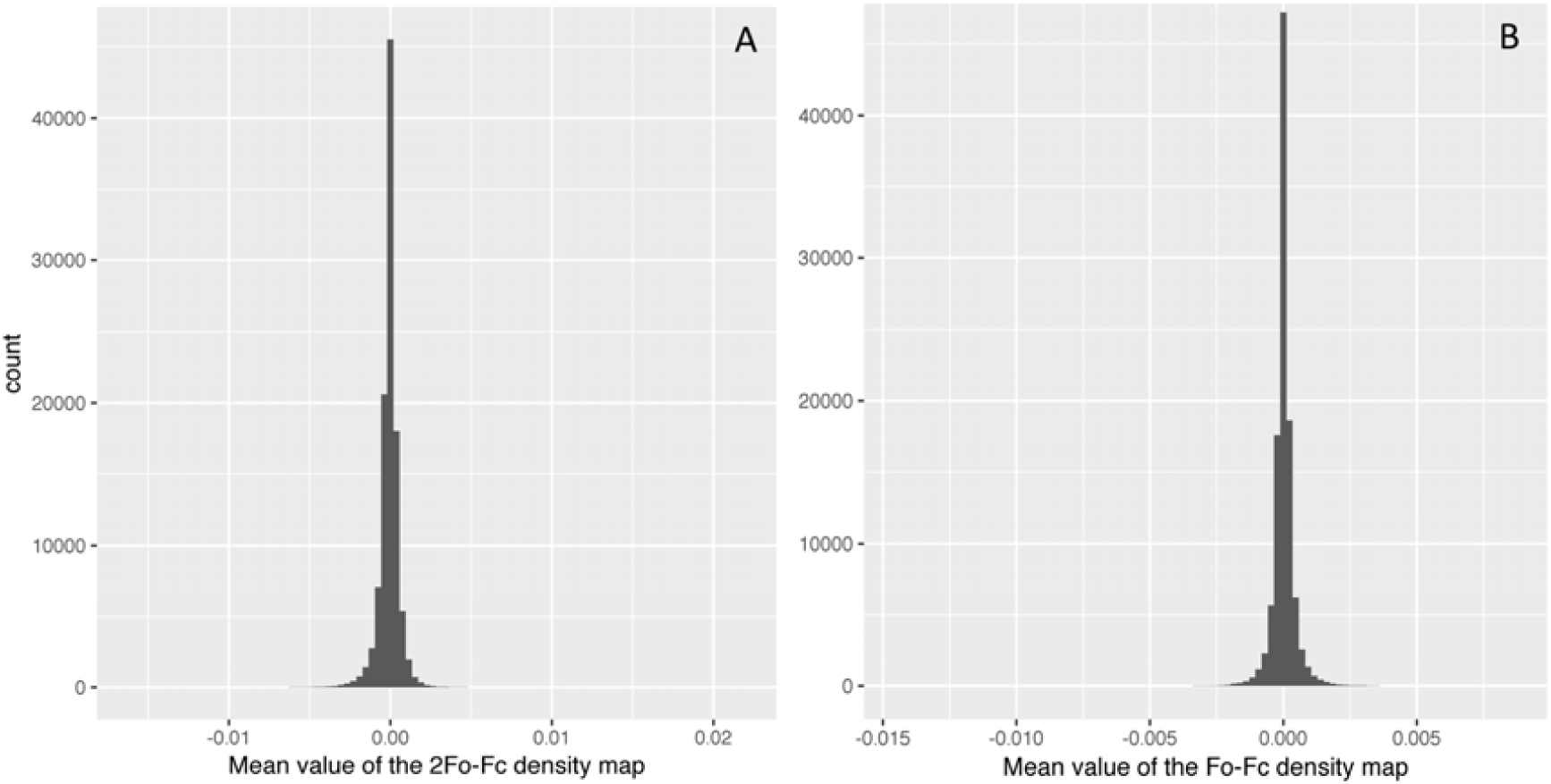
Histogram of the mean value of the 2mFo-DFc map. The histogram illustrates that most of the electron density maps in the PDB are effectively zero-meaned. A) 2Fo-Fc density map, B) Fo-Fc density map.

**Fig 9.**
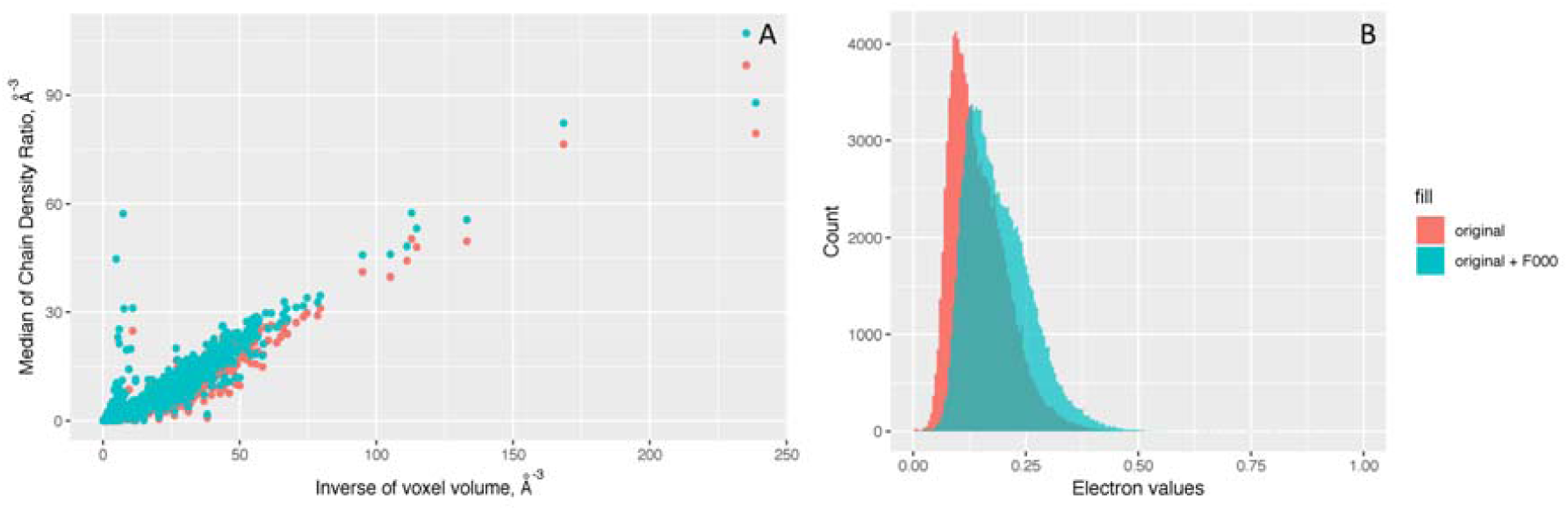
The absolute scales of density ratio for all structures in the PDB. Panel A shows the density ratios vs. inverse of voxel volume plot indicates that there is a consistent 1:3 ratio. Panel B is the histogram of the multiplication of the x and y axes values from Panel A. They both show that the density ratio is not affected much by adding an estimated F_000_ term.

### Evaluative use-case for the electron density conversion factor

One of the most important applications of this work is to estimate the difference density map in terms of electrons. The total difference in expected vs actual electron density can be represented in electron units by dividing the total electron densities by the conversion factor (the median of chain density ratios). As shown in Fig 10, the Fo-Fc map overlaying the 2Fo-Fc map and structure model show several positive (green) and negative (red) density blobs between the measured density from the experiment and the density explained by the given model. On panel A, most of the discrepancies are below six electrons, which can be reasonably interpreted as random background or water noise. Whereas on panel B, there are some difference density blobs worth about 16 and 29 electrons, which could imply actual missing atoms from the model. Moreover, the red missing electron density and the green extra density suggest that the side chain of A389 glutamine should be modeled at the green mesh position rather than the current position. In a similar manner, regions of interest that are common to many PDB entries can be automatically filtered based on electron deviation quality before systematic analysis.

**Fig 10.**
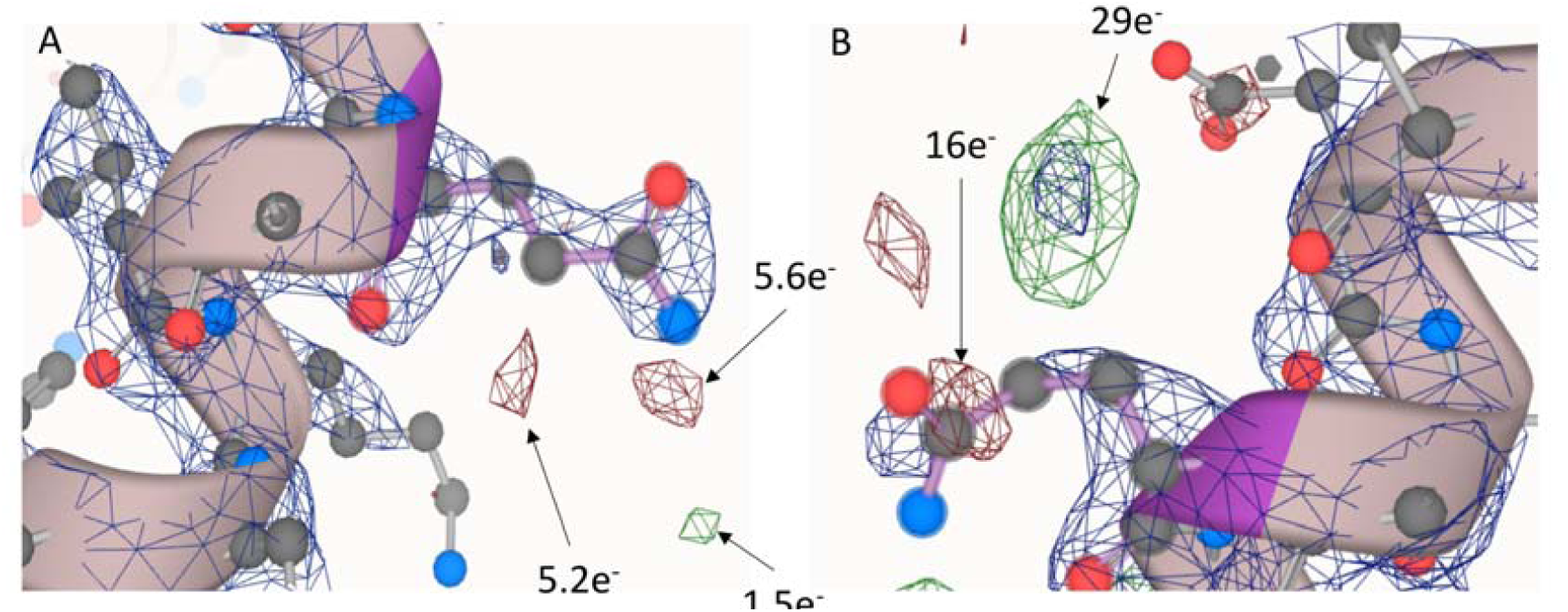
Evaluative use-case for the electron density conversion factor. PDB id: 2P7Z, panel A highlighted residue: A.351, panel B highlighted residue: A.389.

## Discussion

One of the biggest challenges in using electron density maps is that they are in arbitrary scale with no direct physicochemical meaning. This is partly because of missing the magnitude of the structure factor F_000_, which is generally not needed or reported in structure factor files or electron density maps, as it does not affect standard modeling and visualization procedures. Therefore, to put everything back onto an absolute scale so that it is more meaningful for general scientists, this issue needs to be addressed. An approximation of the F_000_ term can be derived from the structure model; however, it is still incomplete because of unknown solvent region compositions and potentially other factors. Also, the zero-mean conversion methods for creation of electron density maps appear to complicate a simple F_000_ correction. This study thus derived new methods that use the median of chain density ratio as a conversion factor to allow the calculation of the missing or excessing electron densities in terms of absolute unit of electrons. These methods implemented in the pdb-eda package provide consistent measures across structures in the PDB with publicly available electron density maps. These new measures are useful for region-specific model evaluation and are suitable for systematic quality control analyses across large numbers of PDB structure entries.

These new methods for deriving a conversion of electron density into a quantity of electrons appear robust with respect to resolution and other PDB entry-specific issues. However, these methods are currently limited to PDB entries containing a significant peptide/protein component. However, the pdb-eda package contains the basic facilities necessary for deriving atomic radii for other polymeric and repetitive supermacromolecular structures. Also, Fig. S1 illustrates a correlation between chain density ratios and resolution, which illuminates a clear path for improvement of atom radii based on resolution.

Over time, the user-base of the wwPDB has shifted from mainly protein crystallographers to a broader community of biologists, computational biochemists, and bioinformaticians, which poses new challenges for how structural data is effectively utilized. While crystallographers are familiar with the concept that not all regions in a structure are of the same quality, this concept is relatively unfamiliar to the other scientists, who tend to focus on global metrics of structure quality like resolution, R-factor, and R-free. Moreover, the experimental details are rather overwhelming for non-crystallographers without extensive training. Thus, this study takes advantages of the recent addition of electron density maps to the PDBe, enabling general scientists to better utilize electron density information now available from the public repository. Our Python pdb-eda package provides easy-to-use methods for interpreting and evaluating structural data with a better physiochemical context. The primary goal of this package is to facilitate a shift in x-ray crystallographic structure evaluation from an entry-specific perspective to a region-specific perspective for the broader scientific community that utilizes the PDB.

## Acknowledgements

This work was supported in part by grants NSF 1252893 (Moseley) and NIH UL1TR001998-01 (Kern). We would like to acknowledge helpful conversations with Dr. David Rodgers.

## Supporting information captions

**Table S1.** Atom type mapping and the electron counts for the 20 common residues.

| Residue | Atom types |
| --- | --- |
| GLY | N: N_single_bb, CA: C_single_bb, C: C_double_bb, O: O_double_bb, OXT: O_intermediate |
| ALA | N: N_single_bb, CA: C_single_bb, C: C_double_bb, O: O_double_bb, CB: C_single, OXT: O_intermediate |
| VAL | N: N_single_bb, CA: C_single_bb, C: C_double_bb, O: O_double_bb, CB: C_single, CG1: C_single, CG2: C_single, OXT: O_intermediate |
| LEU | N: N_single_bb, CA: C_single_bb, C: C_double_bb, O: O_double_bb, CB: C_single, CG: C_single, CD1: C_single, CD2: C_single, OXT: O_intermediate |
| ILE | N: N_single_bb, CA: C_single_bb, C: C_double_bb, O: O_double_bb, CB: C_single, CG1: C_single, CG2: C_single, CD1: C_single, OXT: O_intermediate |
| MET | N: N_single_bb, CA: C_single_bb, C: C_double_bb, O: O_double_bb, CB: C_single, CG: C_single, SD: S_single, CE: C_single, OXT: O_intermediate |
| PHE | N: N_single_bb, CA: C_single_bb, C: C_double_bb, O: O_double_bb, CB: C_single, CG: C_intermediate, CD1: C_intermediate, CD2: C_intermediate, CE1: C_intermediate, CE2: C_intermediate, CZ: C_intermediate, OXT: O_intermediate |
| TRP | N: N_single_bb, CA: C_single_bb, C: C_double_bb, O: O_double_bb, CB: C_single, CG: C_intermediate, CD1: C_intermediate, CD2: C_intermediate, NE1: N_intermediate, CE2: C_intermediate, CE3: C_intermediate, CZ2: C_intermediate, CZ3: C_intermediate, CH2: C_intermediate, OXT: O_intermediate |
| PRO | N: N_single_bb, CA: C_single_bb, C: C_double_bb, O: O_double_bb, CB: C_single, CG: C_single, CD: C_single, OXT: O_intermediate |
| SER | N: N_single_bb, CA: C_single_bb, C: C_double_bb, O: O_double_bb, CB: C_single, OG: O_single, OXT: O_intermediate |
| THR | N: N_single_bb, CA: C_single_bb, C: C_double_bb, O: O_double_bb, CB: C_single, OG1: O_single, CG2: C_single, OXT: O_intermediate |
| CYS | N: N_single_bb, CA: C_single_bb, C: C_double_bb, O: O_double_bb, CB: C_single, SG: S_single, OXT: O_intermediate |
| TYR | N: N_single_bb, CA: C_single_bb, C: C_double_bb, O: O_double_bb, CB: C_single, CG: C_intermediate, CD1: C_intermediate, CD2: C_intermediate, CE1: C_intermediate, CE2: C_intermediate, CZ: C_intermediate, OH: O_single, OXT: O_intermediate |
| ASN | N: N_single_bb, CA: C_single_bb, C: C_double_bb, O: O_double_bb, CB: C_single, CG: C_double, OD1: O_double, ND2: N_single, OXT: O_intermediate, |
| GLN | N: N_single_bb, CA: C_single_bb, C: C_double_bb, O: O_double_bb, CB: C_single, CG: C_single, CD: C_double, OE1: O_double, NE2: N_single, OXT: O_intermediate |
| ASP | N: N_single_bb, CA: C_single_bb, C: C_double_bb, O: O_double_bb, CB: C_single, CG: C_double, OD1: O_intermediate, OD2: O_intermediate, OXT: O_intermediate |
| GLU | N: N_single_bb, CA: C_single_bb, C: C_double_bb, O: O_double_bb, CB: C_single, CG: C_single, CD: C_double, OE1: O_intermediate, OE2: |
|  | O_intermediate, OXT: O_intermediate |
| LYS | N: N_single_bb, CA: C_single_bb, C: C_double_bb, O: O_double_bb, CB: C_single, CG: C_single, CD: C_single, CE: C_single, NZ: N_single, OXT: O_intermediate |
| ARG | N: N_single_bb, CA: C_single_bb, C: C_double_bb, O: O_double_bb, CB: C_single, CG: C_single, CD: C_single, NE: N_intermediate, CZ: C_double, NH1: N_intermediate, NH2: N_intermediate, OXT: O_intermediate |
| HIS | N: N_single_bb, CA: C_single_bb, C: C_double_bb, O: O_double_bb, CB: C_single, CG: C_intermediate, ND1: N_intermediate, CD2: C_intermediate, CE1: C_intermediate, NE2: N_intermediate, OXT: O_intermediate |

**Table S2.**
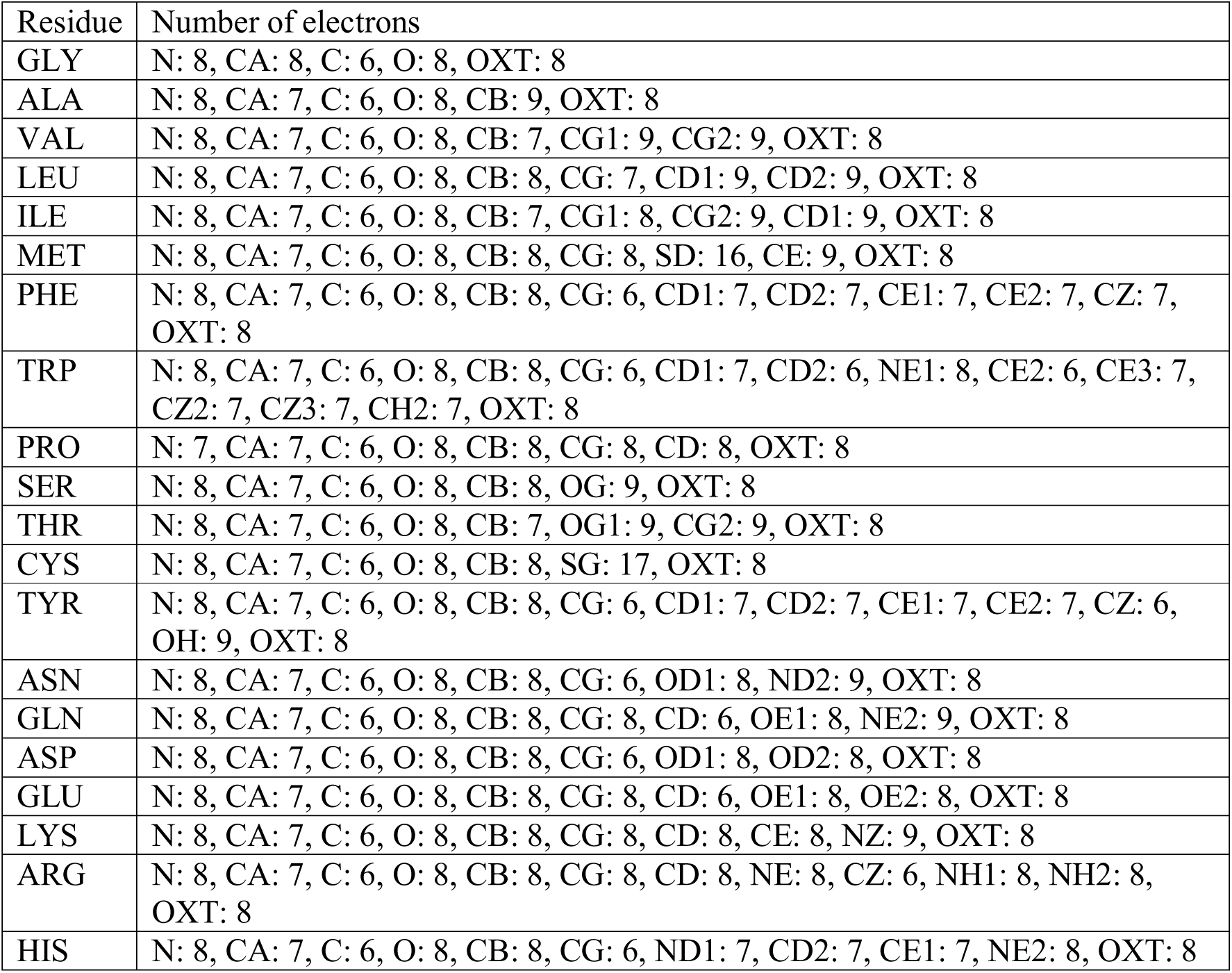
Atom-specific electron counts for the 20 common residues.

**Figure S1.**
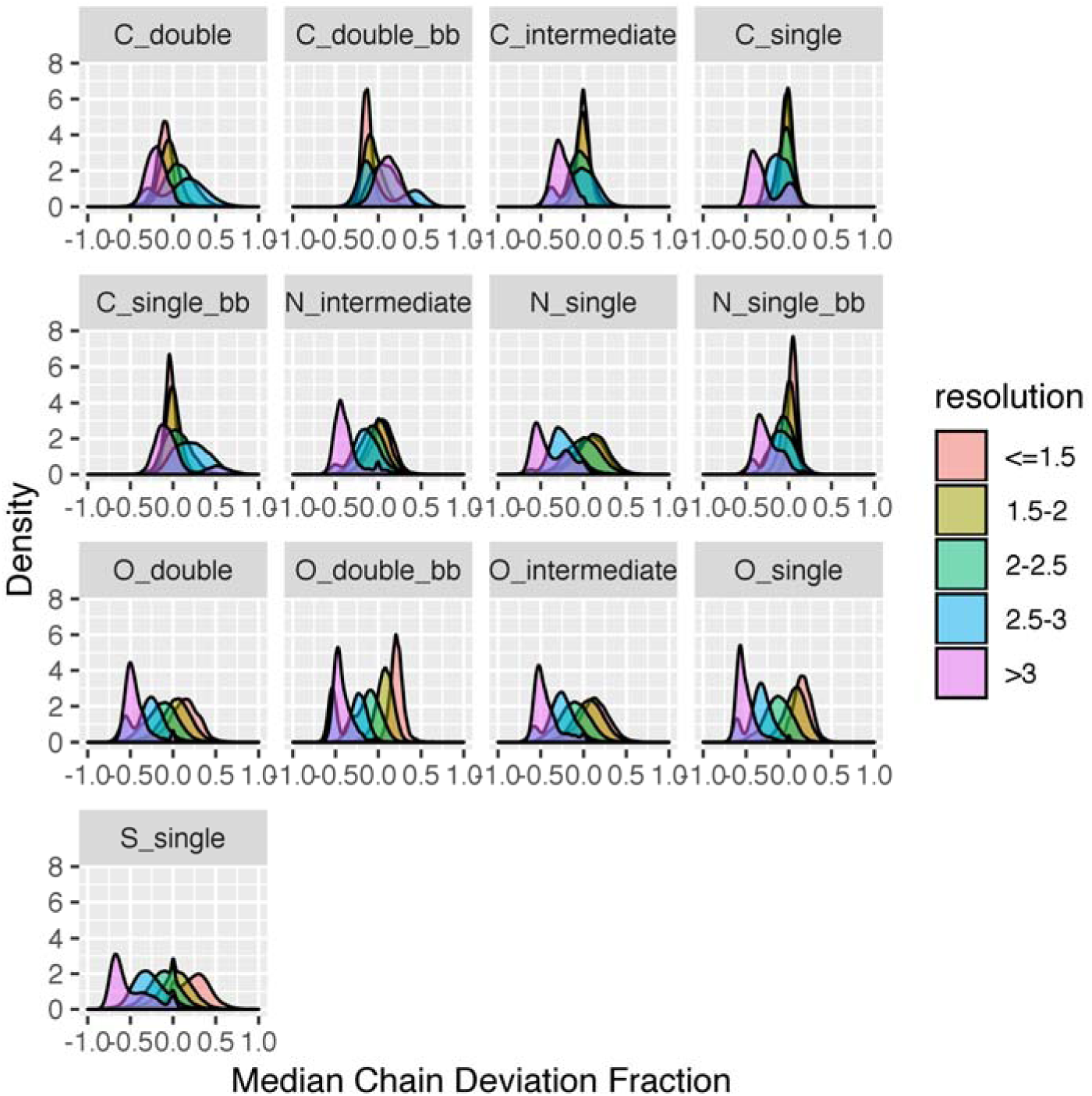
Density plot of the median chain deviation fraction for all structures in the PDB of different resolutions.

## References

1. Berman H, Henrick K, Nakamura H. Announcing the worldwide Protein Data Bank. Nat Struct Biol. 2003;10(12):980.

2. Berman HM, Westbrook J, Feng Z, Gilliland G, Bhat TN, Weissig H, et al. The Protein Data Bank. Nucleic Acids Res. 2000;28(1):235–42.

3. Yao S, Flight RM, Rouchka EC, Moseley HN. A less-biased analysis of metalloproteins reveals novel zinc coordination geometries. Proteins. 2015;83(8):1470–87.

4. Yao S, Flight RM, Rouchka EC, Moseley HN. Aberrant coordination geometries discovered in the most abundant metalloproteins. Proteins. 2017;85(5):885–907.

5. Yao S, Flight RM, Rouchka EC, Moseley HN. Perspectives and expectations in structural bioinformatics of metalloproteins. Proteins. 2017;85(5):938–44.

6. Young JY, Westbrook JD, Feng Z, Sala R, Peisach E, Oldfield TJ, et al. OneDep: Unified wwPDB System for Deposition, Biocuration, and Validation of Macromolecular Structures in the PDB Archive. Structure. 2017;25(3):536–45.

7. Gore S, Sanz Garcia E, Hendrickx PMS, Gutmanas A, Westbrook JD, Yang H, et al. Validation of Structures in the Protein Data Bank. Structure. 2017;25(12):1916–27.

8. Mitroshin I, Garber M, Gabdulkhakov A. Crystallographic analysis of archaeal ribosomal protein L11. Acta Crystallogr F Struct Biol Commun. 2015;71(Pt 8):1083-7.

9. Akey DL, Brown WC, Konwerski JR, Ogata CM, Smith JL. Use of massively multiple merged data for low-resolution S-SAD phasing and refinement of flavivirus NS1. Acta Crystallogr D Biol Crystallogr. 2014;70(Pt 10):2719–29.

10. van Beusekom B, Lutteke T, Joosten RP. Making glycoproteins a little bit sweeter with PDB-REDO. Acta Crystallogr F Struct Biol Commun. 2018;74(Pt 8):463-72.

11. Dutta S, Burkhardt K, Young J, Swaminathan GJ, Matsuura T, Henrick K, et al. Data deposition and annotation at the worldwide protein data bank. Mol Biotechnol. 2009;42(1):1–13.

12. Gutmanas A, Alhroub Y, Battle GM, Berrisford JM, Bochet E, Conroy MJ, et al. PDBe: Protein Data Bank in Europe. Nucleic Acids Res. 2014;42(Database issue):D285–91.

13. EMBL-EBI. Available from: http://www.ebi.ac.uk/pdbe/eds.

14. Kleywegt GJ, Harris MR, Zou JY, Taylor TC, Wahlby A, Jones TA. The Uppsala Electron-Density Server. Acta Crystallogr D Biol Crystallogr. 2004;60(Pt 12 Pt 1):2240–9.

15. Sehnal D, Deshpande M, Varekova RS, Mir S, Berka K, Midlik A, et al. LiteMol suite: interactive web-based visualization of large-scale macromolecular structure data. Nat Methods. 2017;14(12):1121–2.

16. The PyMOL Molecular Graphics System. Version 2.0 Schrödinger, LLC.

17. Read RJ. Improved Fourier coefficients for maps using phases from partial structures with errors. Acta Crystallographica Section A: Foundations of Crystallography. 1986;42(3):140–9.

18. Schwarzenbach D, Abrahams S, Flack H, Gonschorek W, Hahn T, Huml K, et al. Statistical descriptors in crystallography: Report of the IUCr Subcommittee on Statistical Descriptors. Acta Crystallographica Section A: Foundations of Crystallography. 1989;45(1):63–75.

19. Cock PJ, Antao T, Chang JT, Chapman BA, Cox CJ, Dalke A, et al. Biopython: freely available Python tools for computational molecular biology and bioinformatics. Bioinformatics. 2009;25(11):1422–3.

20. Collaborative Computational Project N. The CCP4 suite: programs for protein crystallography. Acta Crystallogr D Biol Crystallogr. 1994;50(Pt 5):760–3.

21. Heyrovska R. Atomic Structures of all the Twenty Essential Amino Acids and a Tripeptide, with Bond Lengths as Sums of Atomic Covalent Radii. arXiv preprint arXiv. 2008;0804(2488).

22. Dorset DL. Filling the missing cone in protein electron crystallography. Microscopy research and technique. 1999;46(2):98–103.

23. Wang B-C. Resolution of phase ambiguity in macromolecular crystallography. Methods in enzymology. 115: Elsevier; 1985. p. 90-112.

